# Novel pink-spot disease in North American kelp nurseries

**DOI:** 10.1101/2025.01.17.633665

**Authors:** Siobhan Schenk, Andrea Sarah Jackman, Jennifer Clark, Laura Wegener Parfrey

## Abstract

Macroalgal cultivation is a growing industry in North America, but is well established in Asia and Africa, where macroalgal disease cause crop losses, placing a significant economic burden on farmers. As kelp cultivation intensifies in North America, kelp disease prevalence is expected to increase in tandem. With input from the kelp-growing community through an online survey, we describe the prevalence of a novel kelp disease, pink-spot disease, which has been observed in Canada and the United States on *Saccharina latissima, Alaria marginata, Nereocystis luetkeana*, and *Macrocystis tenuifolia*. Eight of the fourteen (57%) surveyed kelp growers report bright pink spots (pink-spot disease) on their kelp spools in the land-based nursery stage. Along with the grower survey, we conducted 16S rRNA amplicon sequencing in 2021 and 2022 to investigate the causative agent of pink-spot disease and associated bacterial community changes on infected *Saccharina latissima* (sugar kelp) spools. Our data in both years show that a member of the genus *Algicola* is enriched on visibly diseased spool regions compared to asymptomatic spool regions and may be the causative agent of pink-spot disease. As macroalgal cultivation continues to intensify, monitoring diseases is important to mitigate potential negative impacts.

## Introduction

Macroalgal cultivation is a growing global industry worth 14.7 billion USD in 2019 (FAO, 2021). However, macroalgal cultivation operations (Coleman et al., 2022) and wild macroalgal populations are threatened by direct and indirect effects of climate change (Bindoff et al., 2019; Smith et al., 2024). As macroalgal cultivation increases and climate change continues to intensify, the frequency and severity of macroalgal disease is likely to increase in lockstep (Egan et al., 2013). This pattern has been observed in Africa, where cases of macroalgal disease have increased in prevalence and frequency (Msuya et al., 2022), particularly for macroalgae with low genetic diversity in industrial culture (strain exhaustion), as seen with ice-ice disease of *Kappaphycus* (Ward et al., 2022). Macroalgal disease research has mainly taken place in Asia, where disease costs farmers 15 to 30% of their total crop yields, leading to significant economic losses (Ward et al., 2020). Macroalgal disease appears rare in North America, but this is likely because the industry is comparatively small (1.36% of global production, 0.19 billion USD; FAO, 2021). The existing large-scale negative impacts of disease on macroalgal crops globally, paired with the lack of research into macroalgal disease, poses a risk to the viability and sustainability of the global macroalgal aquaculture industry (Campbell et al., 2019, 2020).

Disease-causing organisms are part of a broad category of host-associated microorganisms known as the microbiota, which includes microeukaryotes, viruses, archaea, and bacteria. Macroalgae-associated bacteria have a strong impact on host development (Marshall et al., 2006; Provasoli & Pintner, 1980), host stress-resistance (Dittami et al., 2016), and host-health (see bacterial infections in Ward et al. (2020)). Due to bacteria’s strong influence on host physiology, researchers have characterized the bacterial community of commercially cultivated macroalgal species, including kelps. This research spans both wild (*Saccharina latissima* (King et al., 2022; Lemay et al., 2018), *Nereocystis luetkeana* (Weigel & Pfister, 2019), *Alaria marginata* (Lemay et al., 2018), and *Macrocystis* sp. (Florez et al., 2017; Lin et al., 2018; Weigel & Pfister, 2019)) and cultivated kelps (*Saccharina latissima* (Davis et al., 2023) and *Alaria* sp. (Davis et al., 2023; Inguanez et al., 2024)). However, research on the microbiota of cultivated kelps is limited (Marzinelli et al., 2024). Understanding the composition of the typical, presumably healthy, microbiota of kelps in cultivation is provides a necessary comparison for the microbiota observed in disease and will enable identification of disease-causing organisms.

Within the seaweed aquaculture sector growers take great care to reduce growth of unwanted organisms (which are sometimes collectively called culture contaminants by growers), including pathogens (e.g., fungus-like oomycetes) and biofouling organisms (e.g. diatoms) (Redmond et al., 2014). Both pathogens and biofouling have the potential to cause significant economic impacts to growers. Some are well described and have relatively effective mitigation techniques, including the addition of germanium dioxide to inhibit biofouling diatom growth and treating *Chondrus spp*. (red macroalgae) cultures infected with oomycetes with sodium dodecyl sulfate (Redmond et al., 2014). However, many unwanted organisms in macroalgal culture are poorly understood and do not have an established preventative or curative protocols. For diseases, one of the main challenges is the difficulty of determining the causative agent of disease.

Traditionally, a microbe’s capacity to cause disease is validated with Koch’s postulates, which requires isolating the disease agent from a diseased organism and infecting a heathy individual with the isolated disease agent, among other requirements (Koch, 1890). Although Koch’s postulates remain relevant today, they are sometimes too stringent (Antonelli & Cutler, 2016) and do not always provide a complete picture of the disease process in marine systems, because marine diseases can occur through multiple mechanisms (Egan & Gardiner, 2016). For example, some diseases are caused by a consortium of microbes (polymicrobial infection; Egan & Gardiner (2016), see coral black band disease; Sato et al. (2016)) and host-associated microbes can switch from an apparently mutualistic or neutral interaction to one that only benefits one of the partners, as seen in the *Phaeobacter gallaeciensis* and *Emiliania huxleyi* system (Egan & Gardiner, 2016; Seyedsayamdost et al., 2011). In addition, visible symptoms may only occur after a preliminary infection or disruption of the microbial community has already taken place (Egan & Gardiner, 2016). In this case, characterizing the microbes associated with visibly diseased regions may describe a community of opportunistic colonizers and the disease agent may no longer be present (Egan & Gardiner, 2016). A lack of clear symptoms early in the disease process is particularly important in the kelp cultivation context, where growers do not have the capacity to thoroughly examine most of their crops closely (e.g. by microscope) due to the scale of biomass production required.

Kelp cultivation can be divided into two main phases, the land-based nursery stage and the open ocean stage (Fig. 1A). In the land-based nursery stage, “kelp seed”—a term used by industry that refers to the development of kelp on spools (twine wrapped around a PVC pipe)—is produced (Fig. 1A). Over approximately six weeks, the kelp develop from spores to juvenile sporophytes (∼2 mm), at which point, the kelp seed are outplanted on the ocean farm, transitioning to the open ocean stage of kelp cultivation. The kelp sporophytes grow in the ocean for four to six months, during which the sporophytes grow in size (∼1 m in length) before being harvested (Fig. 1A).

**Fig 1.**
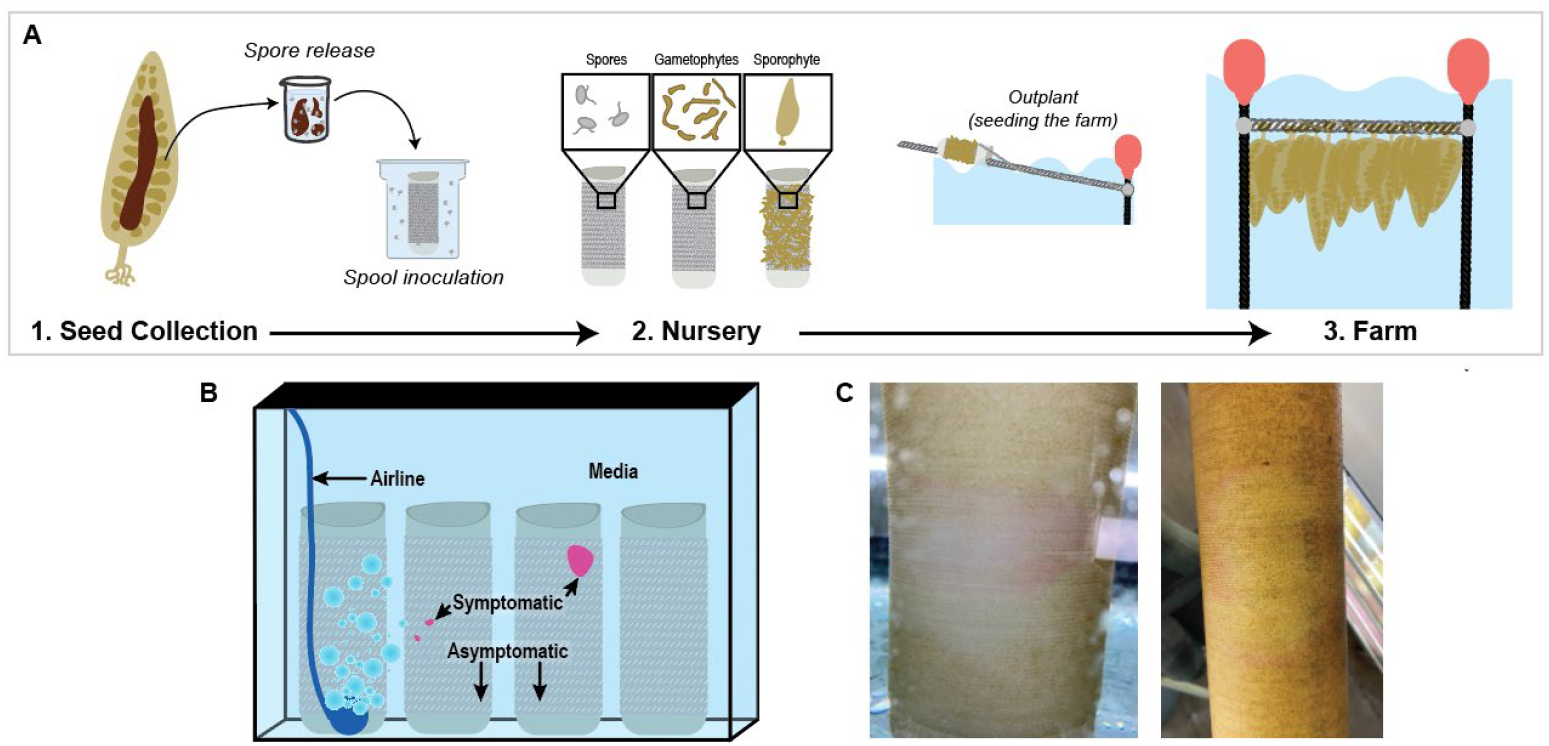
**A)** *S. latissima* lifecycle contextualized within kelp cultivation steps. (1) Kelp seed (wild reproductive sporophytes) is collected and spores are released from the cleaned sori patch. Spores are inoculated onto the spool (twine wrapped around PVC pipes) and maintained in (2) the nursery for approximately six weeks. These kelps spools are the “kelp seed”. (3) The kelp seed is outplanted onto the ocean farm where the sporophytes grow for four to six months before harvest. **B)** Sample type depiction for this study. The symptomatic samples are from visibly pink regions on the spool while asymptomatic samples are from non visibly pink regions. Asymptomatic samples were taken as far away as possible from visibly pink spots. Media and airline samples were also taken to assess the background bacterial community in the aquaria. **C)** Two representative examples of usual presentation of pink-spot disease on *S. latissima* spools in the nursery. The pink spots are less visible in earlier stages of disease and when the spool has a higher density of sporophytes. Note, for images of red-spot disease, see Ezura et al. (1988) figures 1 and 7. Schematics (panel A and B) were designed in Adobe Illustrator

The nursery stage is a cost-intensive and challenging stage in the kelp cultivation process (Coleman et al., 2022). One of the main risks during the nursery stage is biofouling (e.g. diatom blooms) or the development of disease (many reviewed in Ward et al. (2020)). Although there are recent studies examining the bacterial community of diseased kelps (for example, Yan et al. (2023) *Saccharina japonica* malformation disease), many kelp diseases remain either undescribed or were described before the advent of modern sequencing tools (Ward et al., 2020). For this reason, studying the microbial community of kelps at the nursery stage in partnership with growers should be prioritized to establish a better understanding of the healthy nursery microbial community and detect emerging diseases. Doing so may help to safeguard the growing North American macroalgal industry.

Following informal talks with growers, we learned that multiple nurseries in North America have observed bright pink-spots develop on their kelp spools (referred to herein as pink-spot disease). Pink-spot disease is visually similar to red-spot disease previously reported on the kelp *Saccharina japonica* in Japan (Ezura et al., 1988; Sawabe et al., 1998; Yumoto et al., 1989). In red-spot disease, the infection spreads outwards from the origin site along the spool surface as rings, dislodging kelps from spools (Yumoto et al., 1989). Growers observe the same pattern in pink-spot disease. We are not aware of red-spot disease affecting adult kelp sporophytes or non-Laminariales (kelp) algae. These similarities between red-spot and pink-spot disease lead us to hypothesize that they are caused by similar disease agents. Both red-spot and pink-spot disease are a concern for growers because crop loss due to disease causes economic losses. In the case of red-spot and pink-spot diseases, the economic loss is due to visibly disease spool areas becoming bare as the infection progresses. Bare spool regions will not produce harvestable kelp. The prevalence and impact of pink-spot disease is unknown, and to our knowledge, pink-spot disease has not been documented elsewhere in the literature.

We conducted an online survey of kelp growers across North America to 1) assess the prevalence of pink-spot disease, 2) assess the level of concern regarding kelp diseases in general, and 3) start gathering information about disease mitigation strategies already in use by growers. Additionally, we describe the results of an opportunistic sampling of the bacterial community on *S. latissima* spools affected with pink-spot disease. Symptomatic and asymptomatic regions of *S. latissima* spools were sampled from one nursery across two growing seasons. With these data, we aimed to 1) identify a putative pathogen and 2) assess the bacterial community changes associated with pink-spot disease.

## Methods

### Stakeholder survey

We designed a semi-structured survey to quantify the occurrence of pink-spot disease in kelp nurseries and understand the perspective of growers regarding kelp diseases (Table S1). The survey included 17 questions about: disease occurrence, nursery conditions, disease concern, and open-ended questions for participants to describe other evidence of disease, suggest why disease may occur, and provide additional information they felt was important (Table S1). The survey was divided into four sections: (1) contact information (Q1-Q3), (2) disease occurrence and nursery conditions (Q4-Q11), (3) details of the disease (only to be completed if disease was observed in the past year; Q12-Q16), (4) final comments (Q17). Any of the questions could be left blank by the respondent. The survey was distributed to known kelp growers through email, within kelp grower networks, and posted online on the GreenWave community form. Links to the survey are provided in the data availability section.

Of 14 respondent organizations, 7 are based in Canada, 6 in the United States of America, 1 in the United Kingdom, and 1 did not specify. Of these organizations, four focus on kelp restoration, four on research, and six farm kelp commercially. Four of the respondents did not work in an industrial setting.

### Bacterial sample collection and DNA extraction

We collected bacterial samples from a kelp nursery in British Columbia when the nursery team noticed pink-spot disease on *S. latissima* spools in 2021 and 2022. For each aquarium where pink-spot disease was found, we swabbed the symptomatic region (pink; Fig. 1B) and asymptomatic region (not pink; Fig. 1B). The asymptomatic swab was taken as far away as possible from visibly symptomatic regions. We specifically refer to pink and non-pink spool regions as “symptomatic” and “asymptomatic” respectively in our study because with the available data, it is not possible to determine if the asymptomatic regions are healthy regions (uninfected), or if they represent an early, visually asymptomatic, stage of pink-spot disease.

Swabs of small pink spots (below 5 cm^2^) included the entire surface area of the symptomatic area. For larger symptomatic regions, we focused our swabbing effort on the leading edge of the pink spots except for early samples in 2021, where we swabbed the entire symptomatic area (2 samples). Because the pink spots spread outwards, this region of the pink spot should have a bacterial community representative of earlier infection than the middle of the pink spots, which represent a later infection stage. The two whole symptomatic area samples do not appear to be significantly different from the samples where only the leading edge was swabbed, so we include them in our analysis.

We also collected surface swabs of the airline feeding the bubbler (abiotic substrate, Fig. 1), and media samples (F/2 + Germanium dioxide at 8.95 × 10^−3^ g/L; Fig. 1). Surfaces were gently rinsed with filtered seawater prior to swabbing vigorously for 10 s. Media samples were collected by passing a total volume of 50 mL through a 0.2 μM filter (Millipore Sigma, SVGP010) with a pre-rinsed syringe.

In 2021, all samples were taken from a single farm (Farm 1), while in 2022, samples were taken from three farms (Farm 2, Farm 3, and Farm 4). In this context, one farm represents one cycle of spore release from wild-collected sori that will be outplanted in the ocean within 50 km of where the wild sori were collected. For all farms, the kelps are reared in the same nursery for 6 weeks before outplanting, meaning the aquaria have the same water source for all farms in the nursery stage. Water does not freely flow between aquaria, so the spools in different aquaria were physically separated. Aquaria are re-used between farms but are manually scrubbed with 1% bleach and rinsed multiple times with fresh water. Spools from different farms are never combined in the same aquarium.

Bacterial samples were stored at -65°C until transport back to the University of British Columbia on ice, where they were stored at -70°C until DNA extraction with the DNeasy PowerSoil Pro Kit 96-well plate (QIAGEN 47017) in 2021 and the ZymoBIOMICS MagBead DNA/RNA extraction kit (Zymo R2136 and lysis racks S6002-96-3) in 2022. We included one extraction blank per DNA extraction plate both years.

### PCR

The bacterial community samples were profiled using 16S rRNA amplicon sequencing. PCR reactions contained 3 μL of DNA, 15 μL of Phusion Flash High-Fidelity PCR Master Mix (F548L, ThermoFisher), 0.6 μL DMSO, and 2.4 μL of forward (515F, 5’-GTGYCAGCMGCCGCGGTAA-3’) and reverse (806R, 5’-GGACTACNVGGGTWTCTAAT-3’) dual-indexed primers from a 2.5 μM stock, and water to a 30 μL final reaction volume. All PCR reactions consisted of an initial denaturation at (30s at 98°C), 25 cycles of amplification (30 s at 98°C, 30 s at 55°C, and 20 s at 72°C), and a final elongation step (10 min at 72°C). Samples that failed with 25 cycles twice were conducted with 35 cycles, indicated in the metadata. We included one PCR blank per plate.

PCR reactions were quantified with the Quan-IT PicoGreen assay kit (Thermo Fisher, P7589), pooled to equal concentration, cleaned with the PCRs with the Invitrogen PureLink Quick PCR purification kit (K310001), and sent to the University of British Columbia Sequencing Facility for Bioanalyzer quality analysis and Illumina Next Generation Sequencing (MS-1023003 kit; MiSeq v3, 2×300).

### Illumina data processing

Raw, demultiplexed reads were imported into R (v4.4.1; R Core Team, 2022) and processed with the dada2 pipeline (v1.32.0; Callahan et al., 2016). Briefly, forwards and reverse reads were trimmed to 200 bp and primers were removed with the filterandtrim function. Then, amplicon sequence variants (ASVs) were merged into a single sequence table with the mergeSequenceTable function before assigning taxonomy with Silva v138 (McLaren, 2020) formatted for dada2 (McLaren & Callahan, 2021). We removed taxa assigned as chloroplast, mitochondria, eukaryotes, or unassigned at the phylum level. Then, samples with less than 1000 reads, ASVs representing less than 0.001% of the dataset, or found in less than 3 samples were removed. Then, occurrence counts of 3 or less were converted to 0 for non-rarefied data only. Our filtering retained 1,137,331/1,403,759 paired end reads, 767/1,535 ASVs, with a per-sample mean read depth of 27,079.31.

We rarefied our data with coverage-based rarefaction (Chao & Chiu, 2016) with the iNEXT package (v3.0.0; Hsieh et al., 2022) and the metagMisc package (v0.0.4; Mikryukov, 2022) with the coverage set to 0.8 and the rarefaction was iterated 1000 times.

One of our libraries in 2022 included the addition of chloroplast blockers in the PCR reactions (at 1.6 μM) because some samples had a very high number of reads assigned to chloroplasts (blocker information included in associated metadata). Adding the chloroplast blockers decreased the number of chloroplast sequences in the reads by roughly 25% for the samples that included kelp, but did not have an effect on the number of chloroplast sequences in the airline and media samples. We used the general chloroplast blocker 5’-GGCTCAACCCTGGACAG-3’ from (Lundberg et al., 2013) and designed a custom *S. latissima* chloroplast blocker 5’-KKAAATGTAATAGAAACTAC-3’, both ordered from PNA Bio.

To assess consistency in sequencing for the same swab, we performed duplicate PCR reactions for 6 samples (4 symptomatic and 2 asymptomatic) in 2022. The bacterial composition of the reads was extremely similar for duplicate samples (validated visually with a taxaplot and NMDS), so we randomly removed one of the two duplicate samples for data analysis.

### Illumina data analysis

To identify differentially enriched sequences between the symptomatic and asymptomatic regions, we ran a DESeq analysis with the package DESeq2 (v1.44.0; Love et al., 2014) on unrarefied data at the ASV level to retain the highest possible community-level resolution. We used the Wald method with a parametric type fit and set our alpha value to 0.01. Taxa that met the alpha cut off also had to be 3 or more spool samples (total of symptomatic + asymptomatic) to be considered for further analysis. We analyzed both sampling years together because a causative agent should be present across all symptomatic samples. Airline and media samples were excluded from this analysis to allow ASVs that are less substrate specific to be detected by DESeq.

To test for community-wide changes between asymptomatic and asymptomatic spool communities, we ran a PERMANOVA followed by a betadispersion test with the R package vegan (v2.6-8; Oksanen et al., 2022). We also used the package vegan to calculate the Shannon-Weiner diversity index (diversity function). We compared symptomatic and asymptomatic spool sample diversity with an ANOVA. Assumptions of equal sample size and normality were assessed visually with plots, while equal variance was tested with the LeveneTest function in the car package (v3.1-2; Fox & Weisberg, 2019). We ran the PERMANOVA, diversity, and dissimilarity analyses within the same year because different extraction kits were used between years (QIAGEN in 2021, Zymo in 2022), which could influence community composition.

### Tree construction

We used the EukRef pipeline (Campo et al., 2018) to pull closely related ASVs into our tree with a modification for vsearch. First we pulled *Pseudoaltermonas* sequences from Silva v138 (McLaren, 2020) with seqkit (Shen et al., 2016, 2024) and standardized them by converting all uracil (U) bases to thymine (T). We separated the *Algicola* sequences from our ASVs from the rest of the data to prevent clustering and added them back in after the outgroup sequences. We used vsearch to remove sequences less than 500 bp long to improve alignments (Rognes et al., 2016). We manually selected outgroup sequences from a sister clade and added sequences from taxa associated with kelp diseases from Silva v138. All sequences were aligned with mafft (Katoh & Standley, 2013) and trimAl (Capella-Gutiérrez et al., 2009) was used to trim the tree. We removed the following sequences due to exceptionally long branch lengths: AB536964.1.1284, HF912441.1.1438, GU056801.1.1446, and KR054340.1.1206. We generated the backbone tree with RAxML (Stamatakis, 2014) using the GRCAT model with 25 rate categories and 100 bootstrap replicates. We then aligned our ASVs using mothur (Schloss et al., 2009) and placed them in the tree with EPA-ng (Barbera et al., 2019). The final tree was converted to newick format with gappa (Czech et al., 2020) and visualized in figtree (http://tree.bio.ed.ac.uk/software/figtree/).

### Analysis of existing datasets

To assess the prevalence and relative abundance of the putative pathogen from this study, we analyzed 5 other kelp datasets. One of these datasets is of cultivated kelp (Davis et al., 2023), while the others are of wild kelp in British Columbia (Lemay et al., 2018; Park et al. (unpublished); Schenk et al. (unpublished)) and in the United Kingdom (King et al., 2022). All datasets were processed with the dada2 pipeline as described above. We did not filter these data to maximize the chance of detecting our ASVs of interest, even if they are very rare in these other datasets.

## Results and discussion

### Visual description of the pink spots

The pink spots first appear as small patches. The pink is flush along the twine surface and is very difficult to remove fully from the spool. As the pink spots grow, they expand outwards along the spool surface (Fig. 1C). As the infection spreads outwards, the older infection sites become bare of visible kelp (gametophytes or sporophytes). Because we observed pink-spot disease in an active kelp nursery, symptomatic spools were immediately removed from the aquaria and treated with ethanol or bleach (described below) in accordance with the nursery’s disease containment protocol. Thus, the typical growth-rate of the pink spots and the rate of spread to other spools was not studied. However, we noted that asymptomatic spools were present alongside symptomatic spools in the same aquaria when pink-spot disease was detected early (small spots) these asymptomatic spools did not develop pink spots. Removing the symptomatic spools from the aquaria appears to mitigate the spread to other spools to some extent, although we did not quantify this. The majority of aquaria in the nursery remained asymptomatic throughout both the 2021 and 2022 production season.

### Prevalence

Out of the 14 respondents who grow kelp, 8 (57.1%) observed pink spots on spools at some point in their nursery. The affected nurseries were on both the west and east coasts of North America. Of the 8 affected nurseries, some grow multiple kelp species, so we report the number of affected nurseries that observed pink spots on a kelp species at any time (numerator) over the total number of affected nurseries who grow that kelp species (denominator): 7/7 *S. latissima*, 3/3 *Nereocystis luetkeana*, 3/4 *Alaria marginata*, and 1/1 *Macrocystis tenuifolia*. This shows that pink-spot disease affects multiple different kelp species. Growers reported that pink spots on all species were generally small in nature, usually not exceeding ∼ 5 cm^2^ in area before detection and treatment. The causative agent(s) of disease may or may not be the same between kelp species.

### Treatment and prevention

Surveyed growers have a consistent practice of removing the infected spools immediately from nursery aquaria. In less severe cases, spools are immediately spot treated with 1% bleach or 70% ethanol and moved to a quarantine aquarium. The infected spools are maintained in the nursery to assess the efficacy of treatments, which is variable. In severe cases (pink spots larger than ∼5 cm and/or very numerous), spools are immediately removed from the nursery, soaked in 10% bleach, and disposed of.

Respondents identified equipment sterilization between cultivation cycles, regular water changes, water circulation, and appropriate crop density as important measures to prevent disease. All growers who encountered disease stated that rapid intervention to remove infected spools is an important measure to reduce disease severity and prevent spread to other spools. This is in line with practices reported in other published literature (Redmond et al., 2014; Ward et al., 2020).

Most growers who responded to the survey were interested in further collaboration with researchers in general, and to learn more about pink-spot disease specifically. Therefore, we encourage other researchers to collaborate with local kelp growers to target region-specific concerns most effectively.

### Stakeholder’s perspective on disease

We asked survey respondents to report their general level of concern about kelp diseases. We also asked about their concern regarding diatom fouling, to gauge their general level of concern about crop health. In general, there is very little apparent relationship between industry stakeholder concern regarding diatom biofouling and kelp disease. Additionally, there was no apparent relationship between observation of pink-spot disease and disease concern, suggesting that individual perspectives in the industry vary greatly and may be due to factors not captured in our survey. However, 11/15 growers surveyed had at least a moderate level of concern regarding kelp diseases, indicating disease is a concern for growers in Canada and the United States.

### Is there a putative disease agent?

We assayed the bacterial community of symptomatic and asymptomatic regions to identify changes in the community composition associated with disease and gain insight into the underlying causes of disease. As pink-spot disease is visually similar to red-spot disease of *S. japonica*, we hypothesized pink-spot disease may be caused by a similar bacterial pathogen. If pink-spot disease is caused by a single bacterial pathogen, we expect a high and consistent enrichment of a single ASV in symptomatic spool samples across years and farms. Alternatively, if pink-spot disease is caused by a polymicrobial infection or disruption of the normal microbiome due to other stressors, we do not expect to see a consistent enrichment of a single bacterium across symptomatic samples.

To statistically test for differential abundance of taxa by spool health status, we ran a DESeq analysis comparing the symptomatic and asymptomatic spool samples (both years together). DESeq2 identified 16/767 ASVs (2%) that were differentially enriched by spool health (Fig. 3, Table S2). Only 1 ASV (*Algicola* ASV80) is present and enriched in all symptomatic samples. ASV80 has the greatest absolute log2 fold-change value (8.199) of all differentially enriched ASVs, corresponding to a 16-fold increase in symptomatic samples compared to asymptomatic samples. This pattern of enrichment points to *Algicola* ASV80 being strongly associated with pink-spot disease and may be the causative agent of pink-spot disease (Fig. 3). Red-spot disease has been associated with several bacteria, including *Algicola bacteriolytica* (Ezura et al., 1988; Sawabe et al., 1998), *Pseudoalteromonas distincta* (Sawabe et al., 2000), or *Alteromonas* sp. (Sawabe et al., 2000; Yumoto et al., 1989). However, the strongest evidence supports that *A. bacteriolytica* is the causative agent of red-spot disease on *S. japonica* (Ezura et al., 1988; Sawabe et al., 1998).

There are other taxa that show differential enrichment in symptomatic and asymptomatic swab samples; however, these changes are less consistent between years (Fig. 2). This suggests that there are other changes occuring in the bacterial community when pink-spots are present in the aquaria, highlighting the importance of collecting data regarding the typical microbial composition of kelp in the nursery.

**Fig 2.**
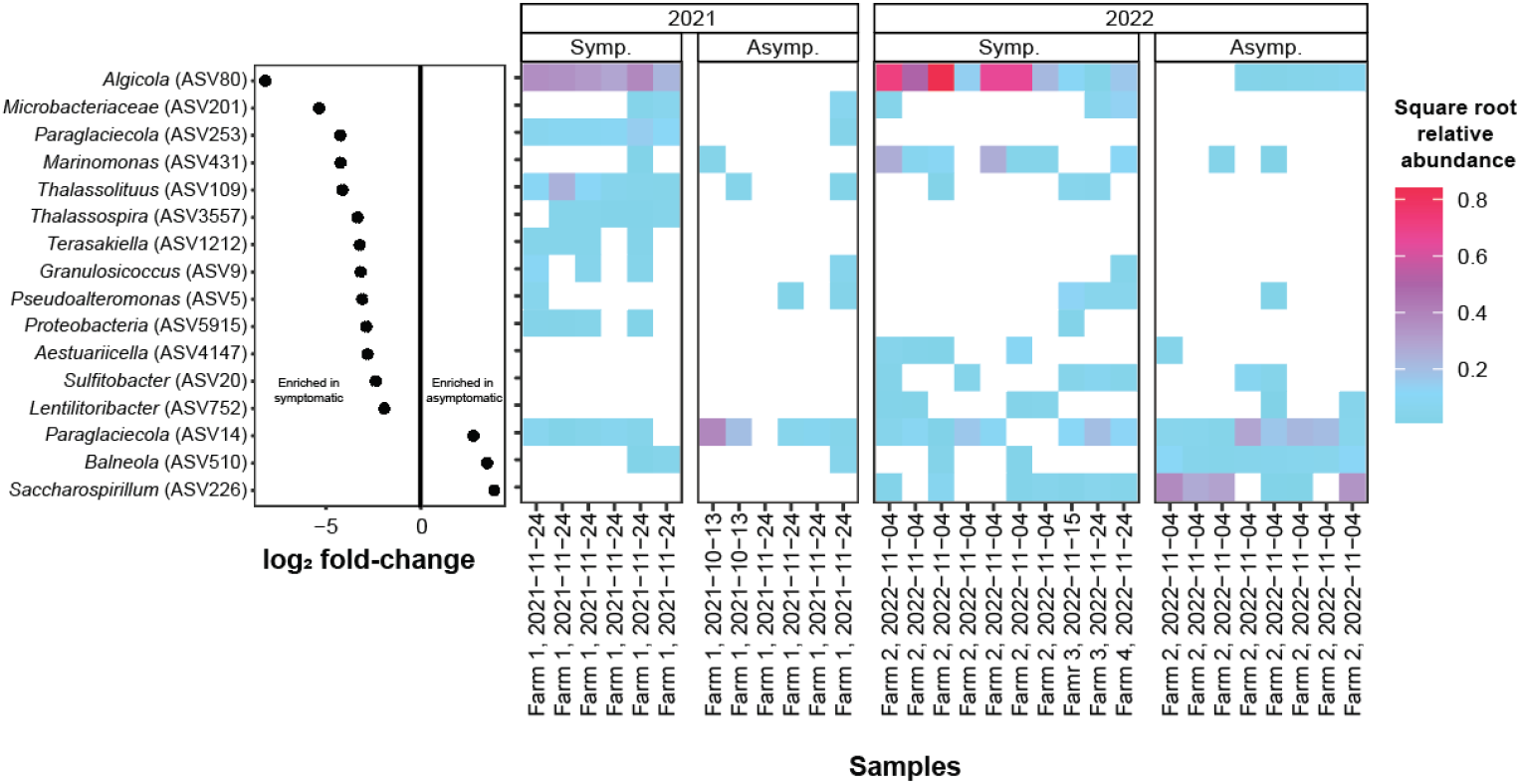
Output of DESeq analysis comparing symptomatic and asymptomatic spools. The left panel y-axis indicates the genus and ASV number that corresponds to the log2 fold-change (x-axis). The right panel shows the square root of the per-sample relative abundance (fill) of the ASV, with the samples divided by spool health status and year (extraction kit). The x-axis shows the sampling date and in 2022, the name of the farm is included. Full DESeq output is available in Table S2

**Fig 3.**
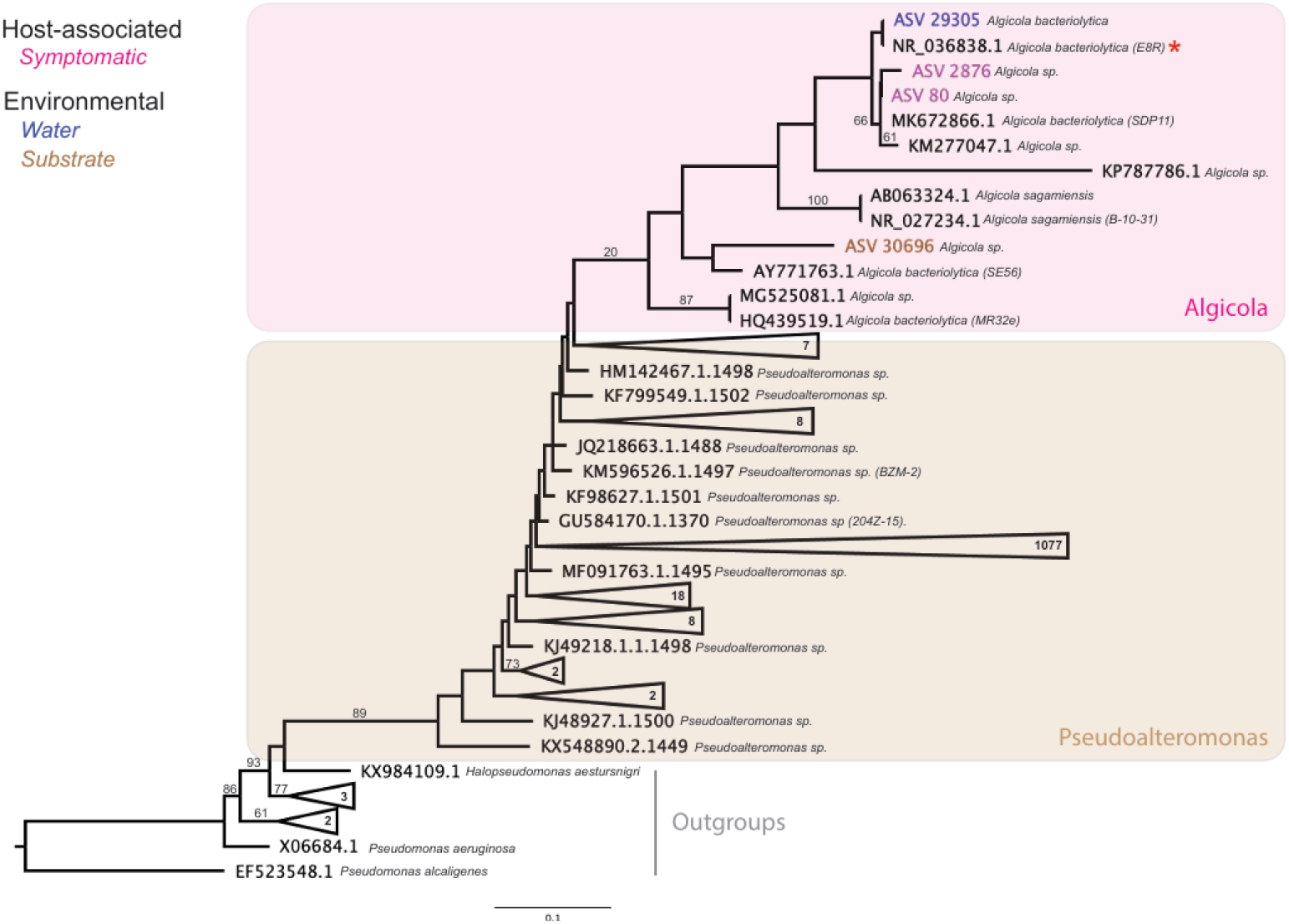
Maximum likelihood (ML) phylogenetic tree of members of [Pseudoalteromonadaceae](https://www.ncbi.nlm.nih.gov/Taxonomy/Browser/wwwtax.cgi?mode=Info&id=267888&lvl=3&lin=f&keep=1&srchmode=1&unlock). This subsection of the tree shows the placement of the genus *Algicola* relative to the Pseudoalteromonas genus. Sequences from Silva are labeled with their accession number, while ASVs from this study and other unpublished studies are labeled with their ASV number corresponding to the taxonomy table and colored by sample type. The proposed pathogen in red-spot disease *A. bacteriolytica* E8R (NR_036838.1) is labeled with an asterix

### Distribution of Algicola in other datasets

We investigated the prevalence of the genus *Algicola* in existing wild and cultivated kelps datasets where pink-spot disease was not observed (Table 1) to assess how commonly *Algicola* is associated with kelp. In our search of multiple unfiltered datasets, we find that the *Algicola* type strain (NR_036838.1) and ASV80 (enriched in pink-spot disease) are not present in any of the datasets, suggesting that these taxa are not ubiquitously associated with kelps.

**Table 1.** Table showing other datasets where we investigated the prevalence of target ASVs. The study with the ENA accession number used to download the reads is listed in the first column, along with the study locale. The next column indicates the sample types and numbers for each study. The following columns show the total number of samples that contained the sequences listed in the column header. The number in the parentheses indicates the mean relative abundance of the ASV in the samples that contained the ASV. Note, we include *Pseudoalteromonas* ASVs in this table as a reference because 1) *Pseudoalteromonas* is closely related to *Algicola* (Fig. 3) and 2) *Pseudoalteromonas* are very fast growing and common in samples, so they contrast the comparatively rare *Algicola*

| Study<br>(ENA accession) | Sample<br>number | *Algalicola* |  |  |  |  | *Pseudoalteromonas* |  |
| --- | --- | --- | --- | --- | --- | --- | --- | --- |
|  |  | NR_036838.1<br>(type strain) | ASV8<br>0 | ASV287<br>6 | ASV28106 | ASV29305 | ASV5 | ASV353 |
| Schenk et al.<br>(unpublished)<br>PRJEB60884<br>British Columbia | kelp (311) | 0 | 0 | 0 | 0 | 0 | 186<br>(0.0151) | 61<br>(0.0006) |
| | water (118) | 0 | 0 | 0 | 0 | 2<br>($<0.0001$) | 89<br>(0.0236) | 68<br>(0.0035) |
|  | substrate (106) | 0 | 0 | 0 | 0 | 0 | 76<br>(0.0175) | 31<br>(0.0017) |
| Park et al.<br>(unpublished)<br>PRJEB64485<br>British Columbia | kelp (93) | 0 | 0 | 0 | 1<br>($<0.0001$) | 0 | 40<br>(0.0024) | 22<br>(0.0007) |
|  | water (5) | 0 | 0 | 0 | 0 | 0 | 4<br>(0.0054) | 0 |
| | substrate (24) | 0 | 0 | 0 | 0 | 0 | 11<br>(0.0032) | 3<br>($<0.0001$) |
| Davis et al.<br>(2023)<br>PRJEB52544<br>British Columbia<br>and Washington | kelp (87) | 0 | 0 | 0 | 0 | 0 | 33<br>(0.0041) | 33<br>($<0.0001$) |
| | water (23) | 0 | 0 | 0 | 0 | 0 | 6<br>(0.0008) | 1<br>($<0.0001$) |
|  | substrate (30) | 0 | 0 | 0 | 0 | 0 | 12<br>(0.0028) | 2<br>(0.0001) |
| King et al. (2023)<br>PRJEB50679<br>United Kingdom | kelp (59) | 0 | 0 | 0 | 0 | 0 | 55<br>(0.0012) | 0 |
| Lemay et al.<br>(2018)<br>PRJEB15100<br>British Columbia | kelp (31) | 0 | 0 | 0 | 0 | 0 | 24<br>(0.0004) | 0 |
|  | water (14) | 0 | 0 | 0 | 0 | 0 | 10<br>(0.0004) | 0 |
|  | substrate (20) | 0 | 0 | 0 | 0 | 0 | 14<br>(0.0012) | 0 |

Next, we expanded out search to include other *Algicola* ASVs present in the pink-spot disease data and the other datasets (Table 1). Here, we identified two water samples from Schenk et al. (unpublished) and one water sample form Park et al. (unpublished)—all from the same sampling site on different days—that contained trace levels of different *Algicola* ASVs (Table 1). These results suggest that *Algicola* is present at low levels in coastal waters where kelp is present, even when the kelps themselves do not show visible signs of disease.

Finally, we probed online databases for records of *A. bacterioytica*. We searched the Encyclopedia of Life (*Algicola bacteriolytica -* EOL, 2024) and the Ocean Biodiversity Information System (*Algicola bacteriolytica* - OBIS, 2024), both of which contained records of *A. bacteriolytica*. These results confirm that *Algicola* is globally distributed in coastal waters. In addition, the OBIS database contained records of *A. bacteriolytica* associated with the kelp *Ecklonia radiata* in Australia, suggesting that *Algicola* can be detected on wild kelps that appear healthy at the time of sampling.

### Algicola phylogeny

*Algicola bacteriolytica* (NR_036838.1) is the proposed causative agent of red-spot disease on *S. japonica* (Ezura et al., 1988; Sawabe et al., 1998), which has a very similar phenotype to pink-spot disease. The main visual difference between these two diseases is the color. A “prodigiosin-like” (red) color in red-spot disease (Ezura et al., 1988; Sawabe et al., 1998) and a pink color in pink-spot disease.

To assess the relatedness of the proposed causative agents of red-spot and pink-spot disease, we constructed a phylogeny of the polyphyletic clade <u>Pseudoalteromonadaceae</u> with sequences available on NCBI. Within this clade, we find that *Algicola* is monophylectic suggesting that *Algicola* is distinct from other taxa in the <u>Pseudoalteromonadaceae</u> (Fig. 3).

Furthermore, we find that ASV80 and *Algicola bacteriolytica* (NR_036838.1) are on different branch tips within the *Algicola* clade, suggesting that ASV80 and NR_036838.1 are distinct taxa. This points to 1) pink-spot disease and red-spot disease being caused by two distinct members of the genus *Algicola* and 2) the capacity of multiple members of the genus *Algicola* to cause diseases during the kelp nursery stage.

### Is the bacterial community different between asymptomatic and symptomatic spools?

To investigate if the entire community differs by disease status, we ran a PERMANOVA for each sampling year (Fig. 4). We ran this analysis both including and excluding all sequences assigned to the *Algicola* genus. We find that differences in bacterial community composition persists after removing *Algicola*, showing that community-wide presence/absence and relative abundance differences are due to changes beyond *Algicola*, aligning with the DESeq output showing polymicrobial shifts in the community by disease status.

**Fig 4.**
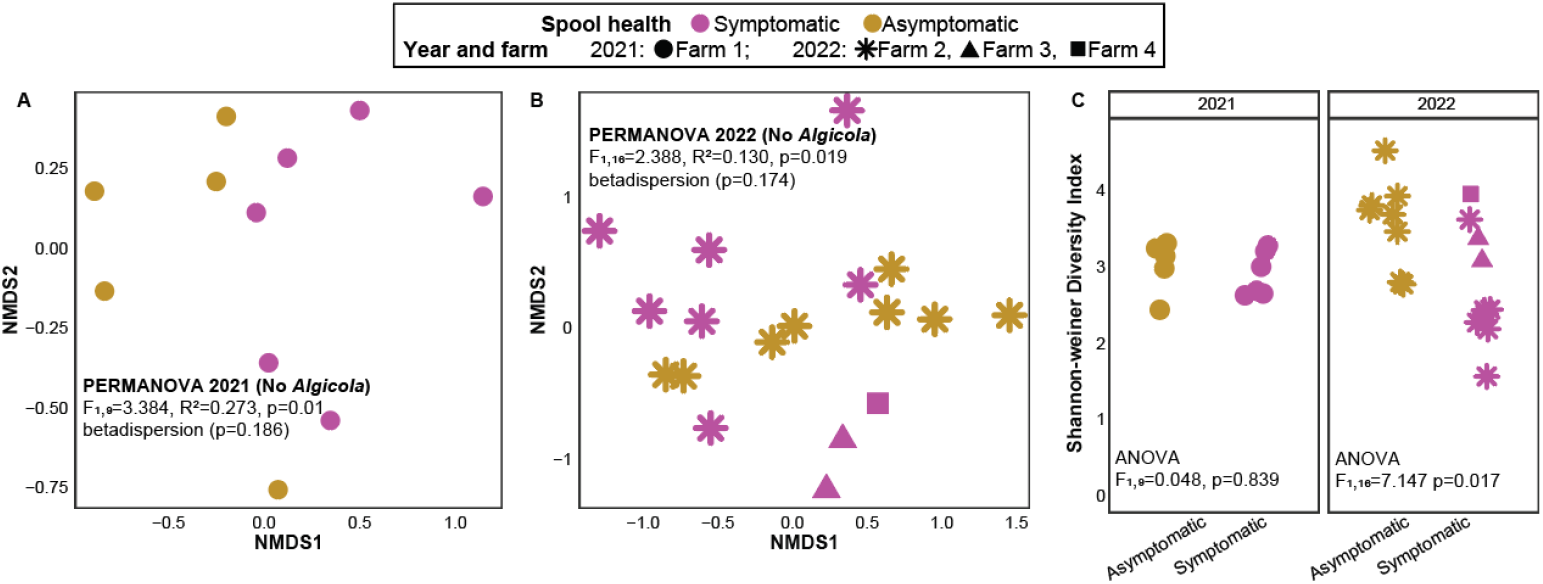
NMDS plot showing spool samples by healthy status with *Algicola* removed from the dataset in A) 2021 and B) 2022. PERMANOVA results are indicated in the corresponding facets along with the p-value of the corresponding betadispersion test. C) The diversity of asymptomatic and symptomatic samples by year (extraction kit)

Although the locale from where sori were collected from 2022 likely has an effect on the bacterial community of the spools, the significant difference between asymptomatic and symptomatic regions persists if we restrict the analysis to only one farm (Farm 2; F_1,14_=2.514, R^2^=0.162, p=0.023). Furthermore, other research (Marzinelli et al., 2015) shows that disease status (bleaching) of the kelp *E. radiata* is more important than spatial variation (analogous to different farms in our data) in explaining variation in the bacterial community. We also tested for differences in alpha-diversity between symptomatic and asymptomatic spool samples (Fig. 4C). In 2022 only, we observe a significant decrease in diversity in the symptomatic samples compared to the asymptomatic samples. It is possible that the inconsistent differences in alpha-diversity between years is due to DNA extraction kits or it could be due to differences in disease progression between years. Future work where pink-spot disease is sampled at multiple time points throughout the disease progression is necessary to reconcile this discrepancy.

Next, to test for higher-level changes in the bacterial community of the spools, we excluded *Algicola* from the datasets and calculated the ten most abundant genera in all four sample types (Fig. 5). We observe consistency in the most abundant taxa (*Colwellia*) by spool health and sampling years, but also some differences between years (*Alteromonas* much more abundant in 2021 and *Sulfitobacter* much more abundant in 2022). Overall, all genera are found in both the asymptomatic and symptomatic samples, showing that there is not a high degree of turnover at the genus level by spool health.

**Fig 5.**
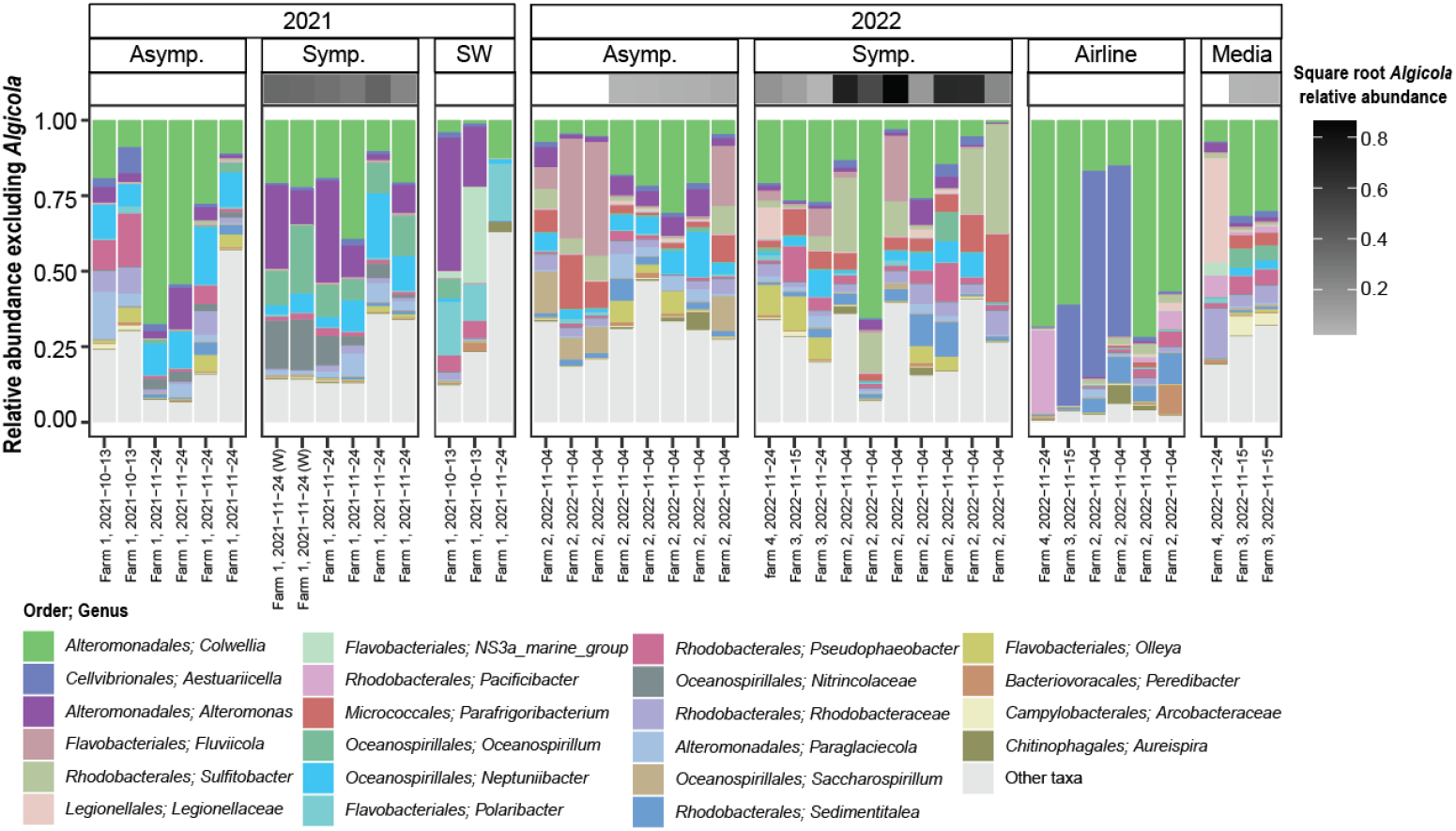
Plot of genera that are one of the 10 most abundant genera by sample type and spool health status excluding the genus *Algicola*. The square root of the relative abundance of *Algicola* is shown in the heatmap on the top of the bars, with white squares showing true 0 relative abundance. The x-axis shows the date the sample was take, along with the farm name in 2022. The facets divide the samples by year (extraction kit), sample type and spool health. Note, for the symptomatic swabs, we indicate the two samples in 2021 where the whole pink area was swabbed (W) in 2021, rather than only the leading edge

## Conclusions

Pink-spot disease is a novel disease found in kelp nurseries and currently present in multiple kelp nurseries across North America. Kelp diseases in the region should be monitored and studied. Growers employ many practices to prevent and mitigate disease in nursery, including sori cleaning with iodine or weak bleach, water filtration, water UV irradiation, and sterilization of culturing equipment in-between uses. Growers also ubiquitously monitor their spools for disease and separate diseased spools from asymptomatic spools to limit the spread of disease.

Our data suggest that pink-spot is a novel disease, likely caused by a member of the genus *Algicola*, as ASV80 *Algicola* was strongly enriched in symptomatic samples and present in all symptomatic samples compared to asymptomatic samples. Pink-spot disease is also accompanied by broader, non-specific changes in the bacterial community, but we cannot determine whether these are a cause or a consequence of disease. ASV80 is closely related but distinct from the causative agent of red-spot disease identified by Sawabe et al. (1998), *A. bacteriolytica*. Isolating ASV80 and testing Koch’s postulates (Koch, 1890) are important next steps to test if ASV80 is the causative agent of pink-spot disease. These efforts should be paired with broader efforts characterizing the nursery microbiota to gain a baseline understanding of what a healthy microbial community in kelp nurseries looks like.

## Declarations

### Funding

Botany Graduate Excellence Award #6372, James Robert Thompson Fellowship, Kruger Graduate Fellowship, Ocean Leaders Fellowship (NSERC) and Internship (NSERC and Cascadia Seaweed) to SS.

Natural Sciences and Engineering Research Council Discovery Grant (RGPIN-2021-03160) and Canada Research Chair tier 2 (CRC-2019-00252) to LWP.

### Competing Interests

Funding for this project was provided in part by Cascadia Seaweed (internship to SS) and JC is employed by Cascadia Seaweed. However, this did not impact the choice of data presented or analyzed in this manuscript.

### Availability of data and material

Raw reads are available at the European Nucleotide Archive (ENA) at PRJEB75651 and the metadata for this study are is available on GitHub siobhanschenk/pink_spot_disease_public

Surveys are available to fill out and view in English https://forms.gle/dsqz7WxcRtQau5D19, Spanish https://forms.gle/2Cc8Pb5s6bifWfZ36, and French https://forms.gle/Ms9qUbpyhbx2EXh6A

### Code availability

Script for this study are available on GitHub siobhanschenk/pink_spot_disease_public

### Authors’ contributions

Study conceptualization and data collection were performed by SS and JC. Data analysis and initial manuscript drafts by SS and AJ. All authors contributed equally to editing and finalizing the manuscript. All authors read and approved the final manuscript.

## Acknowledgements

All respondents who provided their time and valuable perspective by answering our survey.

B. Sedano-Herrera who expanded the potential reach of our research by translating the survey from English to Spanish.

E. Morien whos’ bioinformatics expertise was essential to construct the phylogeny.

Colleagues who provided feedback on the manuscript drafts: N. Salland, P. Lund and V. Supratya.

Additionally, we would like to thank Cascadia Seaweed who provided funding and access to their nursery and farms for this project. In particular, we thank the nursery team: T. Campbell, J. Carson-Austin, L. Krzus, S. Young, and the operations team, S. Martin and C. Bates, for their help setting up the experiments and sampling.

## Supplement

**Table S1.**
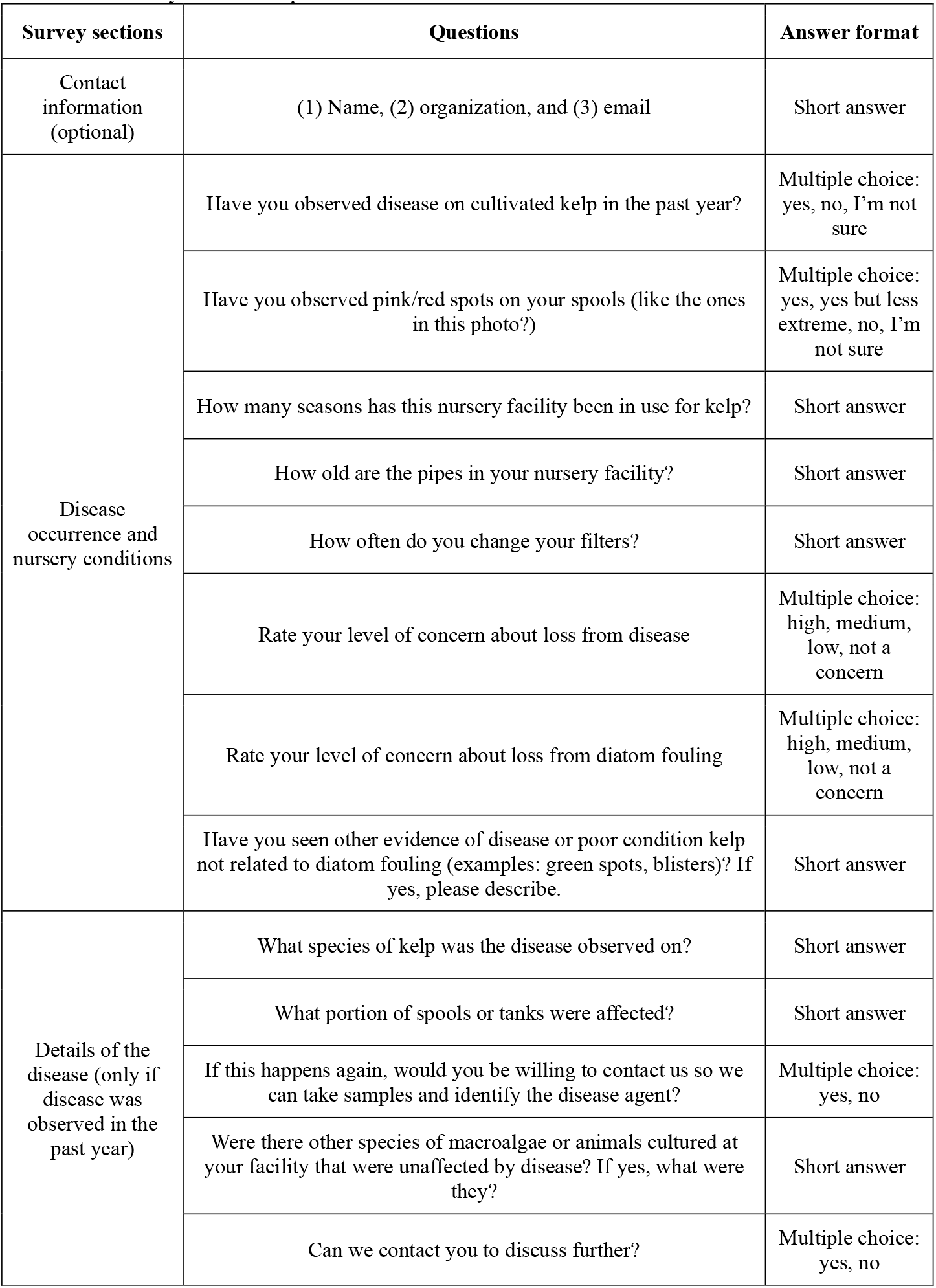
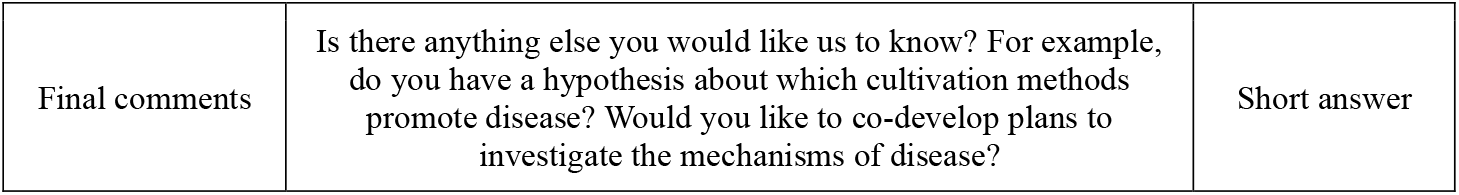
Survey sections, questions, and answer format.

